## Supplemental Materials for "Hypertrophic cardiomyopathy mutations at the folded-back sequestered β-cardiac myosin S1-S2 and S1-S1 interfaces release sequestered heads and increase myosin enzymatic activity"

**Table S1**. Actin-activated ATPase activity values for 2-hep HMM and 25-hep HMM

| | **2-hep HMM $k_{cat}$** | **25-hep HMM $k_{cat}$** | **(25-hep HMM:2-hep HMM)** |
|---|---|---|---|
| **WT** | 2.39 ± 0.06 | 1.38 ± 0.06 | 0.57 ± 0.03 |
| **R249Q** | 1.69 ± 0.08 | 1.63 ± 0.06 | 0.96 ± 0.06 |
| **H251N** | 3.31 ± 0.05 | 2.95 ± 0.07 | 0.91 ± 0.03 |
| **D382Y** | 3.21 ± 0.06 | 2.72 ± 0.12 | 0.84 ± 0.04 |
| **R719W** | 2.68 ± 0.04 | 2.79 ± 0.07 | 1.03 ± 0.03 |
| **D/R** | 1.35 ± 0.02 | 1.41 ± 0.03 | 1.04 ± 0.04 |
| **I457T** | 4.19 ± 0.33 | 2.35 ± 0.31 | 0.56 ± 0.10 |

From the $k_{cat}$ values in table S1, it is clear that H251N 2-hep HMM has a ~40% higher $k_{cat}$ than WT 2-hep HMM, and R719W 2-hep HMM has a ~10% higher $k_{cat}$ than WT 2-hep HMM. These values agree with previously published experiments on sS1 myosin with the same mutation.

**Table S2**. Slow phase fraction (SRX) of 2-hep and 25-hep HMM as measured using a single turnover mant ATPase assay

| | **2-hep** | **25-hep** |
|---|---|---|
| **WT** | 19 ± 3% | 59 ± 7% |
| **R249Q** | 22 ± 1% | 21 ± 3% |
| **H251N** | 19 ± 1% | 35 ± 4% |
| **D382Y** | 18 ± 4% | 32 ± 4% |
| **R719W** | 18 ± 3% | 19 ± 7% |
| **D/R** | 22 ± 9% | 27 ± 5% |
| **I457T** | 18 ± 1% | 44 ± 4% |

**R249Q characterization**
R249Q has been reported as a pathogenic HCM mutation[1]. As seen in Table S1, the actin-activated ATPase rate for R249Q 2-hep HMM is lower than that of the WT 2-hep HMM by 33 ± 4% (we see a similar decrease in ATPase using sS1). We further characterized the mutation using the in vitro motility assay using sS1 and found that the velocity was 50 ± 5% lower. However, we have also shown previously that R249Q weakens the binding between the S1 head and the S2 tail of myosin[2], and the ATPase data in this manuscript shows that R249Q's weakening of the S1-S2 interaction leads to increased activity of 25-hep HMM. These data suggest that even though the mutation may lead to a loss in the activity of individual myosin motors, it releases more myosin heads from the closed state making them available to interact with the actin in the sarcomere, thereby leading to hypercontractility.

**D382Y characterization**
The D382Y mutation results in $k_{cat}$ values for 2-hep HMM that are 33 ± 4% higher compared to the WT 2-hep HMM. Our previous experiments using the in vitro motility assay showed that the velocity of D382Y was 6 ± 2% lower than the WT. In this case, we have a significant gain in function in the actin-activated ATPase rate, a very small decrease in the actin gliding velocity, and a significant decrease in the percentage of heads in the SRX. Overall, these data are consistent with the D382Y mutation causing hypercontractility at the molecular level.

### Supplemental Methods

*Expression and purification of recombinant human β-cardiac myosin constructs*
The human β-cardiac 2-hep HMM cDNA consists of a truncated version of MYH7 (residues 1-855), corresponding to S1 and the first two heptad repeats of S2, followed by a GCN4 leucine zipper[3] to ensure dimerization. This is further linked to a flexible GSG (Gly-Ser-Gly) linker, then an eGFP moiety followed by another GSG linker, and finally ending with an 8-residue (RGSIDTWV) PDZ-binding peptide. The human β-cardiac 25-hep HMM construct was similar to the 2-hep HMM, except that MYH7 was truncated after 25 heptad repeats (175 amino acids) of the S2 region (up to residue 1016).

Recombinant human β-cardiac 2-hep HMM and 25-hep HMM containing HCM-causing mutations were co-expressed with a FLAG-tagged human ventricular cardiac essential light chain (ELC) in C2C12 mouse myoblast cells using adenoviral vectors (pAd Easy system Qbiogene). C2C12 cells were grown at 37°C and 8% $CO_2$ in growth medium (DMEM + 10% fetal bovine serum + 1x pen-strep). 10 plates of confluent C2C12 cells were differentiated to myotubes by adding differentiation medium (DMEM + 2% horse serum + 1x pen-strep). The cells were differentiated for 2 days. Next, the cells were infected with adenoviruses carrying the myosin sS1 and myosin FLAG-ELC in growth media ½ (DMEM + 5% fetal bovine serum + 1x pen-strep). The cells were infected for 4 days and then harvested. The media was removed from the cells, and they were washed with ice cold PBS. Then 1 ml of lysis buffer (20 mM imidazole pH7.5, 100 mM NaCl, 4 mM $MgCl_2$, 1 mM EGTA, 1 mM EDTA, 0.5% tween-20, 1 mM DTT, 3 mM ATP,

1 mM PMSF, 10% sucrose and Roche protease inhibitors) was added to each plate and the cells were harvested by scraping. The cells were lysed using a dounce homogenizer, and the lysate clarified by centrifugation at 23,000 rpm for 20 min using a Ti60 ultracentrifuge rotor. The supernatant was then incubated with 75 µL of anti-FLAG resin per 15 cm dish for 1.5 - 2 hours at 4°C. This allowed the HMM to bind to the anti-FLAG resin.

Next, the mouse RLC was stripped from the protein with 0.5% Triton X-100, 5 mM CDTA, 200 mM KCl and 20 mM Tris pH 7.5 at 4˚C for 75 min. Human RLC binding to the heavy chain was performed by incubation of an excess amount of human cardiac RLC for 2 hr at 4˚C. The protein was eluted from the anti-FLAG resin by incubating it with TEV protease overnight at 4˚C, and it was purified the next day using anion exchange chromatography. Analysis of a 15% SDS PAGE gel confirmed a 1:1:1 stoichiometric complex of HMM heavy chain with both human light chains. The RLC on the human HMM is unphosphorylated. All experiments presented in this manuscript were performed using unphosphorylated HMM.

*Actin activated ATPase assay*

Only freshly prepared HMM was used for ATPase assays. Since we aimed to elucidate whether HCM mutations alter the intramolecular interactions of myosin, we always prepared the 2-hep HMM and 25-hep HMM for each mutation in parallel, and the assays were performed on both proteins simultaneously.

To prepare F-actin, G-actin was dialyzed extensively into ATPase buffer to remove any residual ATP. Actin concentration was then measured using absorbance at 290 nm in a spectrophotometer. The steady-state actin-activated ATPase activities of the WT 2-hep and 25-hep HMM and mutant human β-cardiac 2-hep and 25-hep HMMs were determined using a colorimetric assay to measure inorganic phosphate production at various time points (0 - 30 min) from a solution containing myosin (0.01 mg $ml^{-1}$), ATP and varying concentrations of actin filaments (0 - 100 µM) [4]. All measurements were made at 23°C. The time-dependent rate for each actin concentration was calculated by fitting the phosphate signal as a function of time to a linear function. The slope was then converted to activity units normalized to a single myosin head. Kinetic parameters (i.e. $k_{cat}$) were extracted from the data by fitting the activity at each actin concentration to the Michaelis-Menten equation to determine maximal activity using the curve fitting toolbox in MatLab[5]. The errors in the fitted values were determined using 100 bootstrap iterations.

*Microscale thermophoresis for sS1-S2 binding affinity*

To assess if the mutations at the S1-S1 interface altered the interaction between S1 and S2, we used microscale thermophoresis (MST). To study this interaction, we used freshly prepared 2-hep HMM with its C-terminal eGFP, and unlabeled S2 expressed in and purified from bacteria (amino acids 839-968)[2]. The proximal S2 construct includes the first 126 amino acids of S2 and begins 4 residues before the end of S1. Due to the low affinity between S1 and S2, the S2 fragment was concentrated to >300 µM to get a binding curve. Both proteins were dialyzed into MST assay buffer (10 mM Imidazole pH7.5, 100 mM KCl, 1 mM EDTA, 2 mM $MgCl_2$, 1 mM DTT, 500 µM ADP and 0.05%

tween). Before using them for the assay, both proteins were centrifuged at 100,000 rpm in a TLA 100 rotor for 20 min to remove any aggregates. For the assay, we used 16 serial dilutions of S2 myosin starting at > 300 μM, with a 3-fold dilution for each subsequent sample. The 2-hep HMM was kept constant at 50 nM. The samples were loaded into NT.115 premium treated capillaries and incubated at 23°C for 45 minutes in the dark. All MST data was recorded at 23°C. The 2-hep HMM-S2 interaction was followed by monitoring the eGFP fluorescence. A blue LED at 30% excitation power (BLUE filter; excitation 460-480 nm, emission 515-530 nm) and IR-Laser power at 60% was used. Data analysis was performed with the software NTAffinityAnalysis (Nanotemper Technologies) where the binding isotherms were derived from the raw fluorescence data. The binding isotherms were then fitted using Matlab, with the Hill equation for cooperativity, to estimate an apparent dissociation constant ($K_D$), using a linear regression method. There were preparation-to-preparation differences in the binding affinity, and the WT $K_D$ ranged between 35-50 μM; however, the relative differences in binding affinities between the WT and mutant myosins were constant.

*Single turnover experiments*

Single turnover experiments were performed in a fluorescence plate reader (Tecan model – Infinite M200 PRO). Experiments were performed with the WT and mutant versions of 25-hep HMM and 2-hep HMM constructs of human ß-cardiac myosin as described[6]. Because of the slow rates of the myosin basal ATPase and SRX-based nucleotide release along with difficulties in expressing large quantities of human cardiac myosin, a low volume, plate-based measurement is an apt choice for measuring the single turnover rates. Appropriate controls and comparisons to a stopped-flow measurement were performed by Anderson et al.[6] to ensure that we are not missing any fast phase in the plate-based measurement. Briefly, these experiments were performed in a 96-well plate (Greiner polypropylene microplate) by mixing HMM in a buffer containing 10 mM Tris pH 7.5, 4 mM $MgCl_2$, 1 mM EDTA, 1 mM DTT and 5 mM KAc with 2'-(or-3')-*O*-(*N*-Methylanthraniloyl) adenosine 5'-triphosphate (mant-ATP, Thermo Fischer Scientific) at a final concentration of 100 nM. After 10 s, 2 mM ATP was added, followed by measuring the fluorescence signal at 470 nm after excitation at 405 nm. Fluorescence was recorded every ~2 s for 16 min total and the traces were normalized and plotted[6]. The kinetic traces were fitted to a bi-exponential decay function which yielded the amplitudes and rates of the fast (DRX rate) and slow (SRX rate) phases.

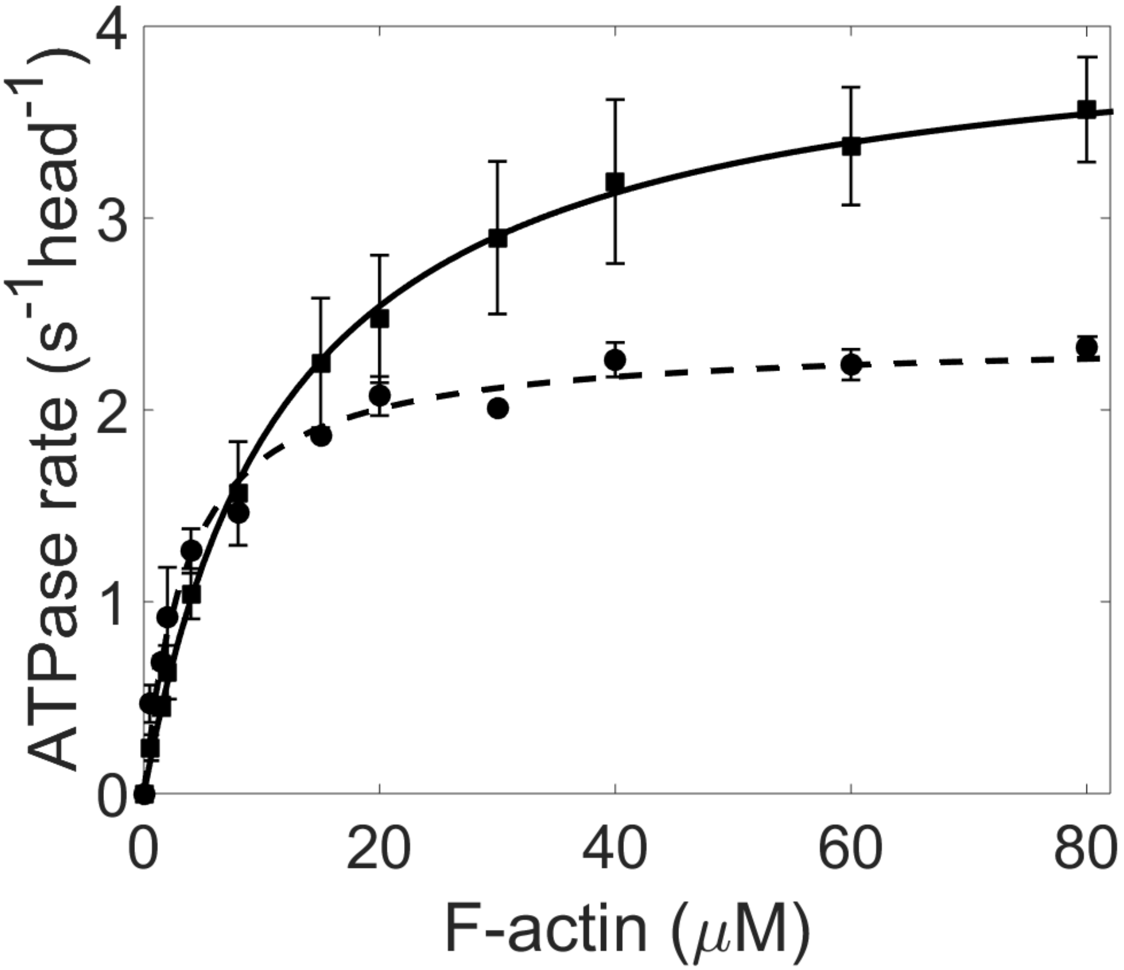


Supplemental Figure 1. Actin-activated ATPase of WT 2-hep HMM (circles and dashed line) vs I457T 2-hep HMM (squares and solid line) clearly shows that I457T has a higher $k_{cat}$ compared to WT 2-hep HMM.

**Supplemental Video 1**: WT 2-hep HMM in-vitro motility (scale bar is 10 µm)
**Supplemental Video 2**: I457T 2-hep HMM in-vitro motility (scale bar is 10 µm)
